# Temperature-dependent competitive outcomes between the fruit flies *Drosophila santomea* and *D. yakuba*

**DOI:** 10.1101/2020.07.25.220863

**Authors:** Aaron A. Comeault, Daniel R. Matute

## Abstract

Changes in temperature associated with climate change can alter species’ distributions, drive adaptive evolution, and, in some cases, cause extinction. Research has tended to focus on the direct effects of temperature, but changes in temperature can also have indirect effects on populations and species. Here we test whether temperature can indirectly affect the fitness of *Drosophila santomea* and *D. yakuba* by altering interspecific competitive outcomes. We show that, when raised in isolation, both *D. santomea* and *D. yakuba* show maximal performance at temperatures near 22°C. However, when raised together, *D. santomea* outcompetes *D. yakuba* at a lower temperature (18°C), while *D. yakuba* outcompetes *D. santomea* at a higher temperature (25°C). We then use a ‘coexistence’ experiment to show that *D. santomea* is rapidly (within 8 generations) extirpated when maintained with *D. yakuba* at 25°C. By contrast, *D. santomea* remains as (or more) abundant than *D. yakuba* over the course of ~10 generations when maintained at 18°C. Our results provide an example of how the thermal environment can indirectly affect interspecific competitive outcomes and suggest that changes in the competitive advantage of species can lead to some species becoming more prone to extinction by competitive exclusion.

## Introduction

Species vary in their physiological tolerance and behavioral preference for different thermal environments (Calosi et al. 2010; Kellermann et al. 2012). Temperature is therefore an important abiotic factor that can drive local adaptation (McNab 1971; Freckleton et al. 2003; Campbell-Staton et al. 2016, 2018; Stager et al. 2016; Delhey 2017, 2019) and shape species’ ranges (Soberón 2007; Calosi et al. 2010; Early and Sax 2011; Kellermann et al. 2012). However, a species’ range and their response to variation in climate is also shaped by biotic interactions. Competition is one outcome of biotic interactions that, like temperature, can drive phenotypic and ecological divergence (i.e. character displacement (Pfennig and Pfennig 2009; Stuart and Losos 2013)), the maintenance of intraspecific variation (Roughgarden 1972; Bolnick 2001; Harris et al. 2008), speciation (Polechová and Barton 2005; Winkelmann et al. 2014), extinction (Park 1954; Davis et al. 1998*a*; Alexander et al. 2015), and global biogeographic patterns (Pianka 1966; Willig et al. 2003). The majority of biotic interactions, such as competition, occur across a range of abiotic conditions (either temporally or geographically). To understand how biotic interactions, such as interspecific competition, affect a species abundance and evolution, it is important to understand the outcome of those interactions across different abiotic conditions (e.g. thermal environments (Davis et al. 1998*a*; Alexander et al. 2015)).

The outcome of interspecific competition has been shown to vary with temperature for a number of species inhabiting different environments. For example, experimental work in communities of algae (Goldman and Ryther 1976; Hillebrand 2011), beetles (Park 1954; Wilson et al. 1984), fungi (Carreiro and Koske 1992), alpine plants (Klanderud and Totland 2007; Alexander et al. 2015), and fruit flies (Davis et al. 1998*a*, 1998*b*) have all shown how temperature can indirectly affect a species’ relative abundance through competition. However, direct effects of climate and competition have also been reported: experimental communities of tussock tundra plant communities show no noticeable interaction between temperature and competition (Hobbie et al. 1999). Therefore, while the majority of studies point towards temperature as an important abiotic control on the fitness consequences of interspecific competition, more examples are required to understand its generality.

One aspect of temperature-mediated competitive outcomes that remains underexplored is its prevalence in closely related (e.g. sibling) species. Species that have diverged recently and come into secondary contact are likely to share more aspects of their ecology than distantly-related species, potentially leading to strong competition (Schluter and McPhail 1992; Grant and Grant 2006). Species pairs that display overlapping or adjacent (parapatric) geographic ranges therefore provide useful systems in which to test for variable outcomes of interspecific competition, especially when those ranges occur along an environmental gradient. The sibling species *Drosophila santomea* and *D. yakuba* provide one such system. *D. santomea* is endemic to the cool tropical forest of the island of São Tomé in the Gulf of Guinea. In laboratory experiments adult *D. santomea* show a behavioural preference for moderate temperatures (22°C) and larval survival and egg hatchability drops at temperatures above 25°C (Matute et al. 2009). *D. yakuba*, on the other hand, is a generalist species that has a broad distribution across sub saharan Africa and is regularly found in association with human-modified habitats (Lachaise et al. 1988; Cooper et al. 2018). In nature, populations of *D. yakuba* therefore experience environments subject to a much wider range of temperatures than populations of *D. santomea*, and *D. yakuba* can tolerate an overlapping but broader range of temperatures in the lab (Matute et al. 2009; Cooper et al. 2018). On the island of São Tomé the distributions of *D. santomea* and *D. yakuba* are adjacent, with *D. santomea* typically occurring at forested habitats above 800 m and *D. yakuba* at lower elevation open habitats (Llopart et al. 2005a; Comeault et al. 2016). The two species form a narrow hybrid zone between 800 and 900m elevation that has been stable for over 20 years and occurs as lowland agricultural fields give way to upland rainforest habitats (Lachaise et al. 2000; Llopart et al. 2005b; Matute 2010a; Comeault et al. 2016). The fact that *D. santomea* and *D. yakuba* show differences in their realized niche in nature, yet display broadly overlapping thermal tolerances in the lab raises the question of how interspecific competition may contribute to the maintenance of their distinct ecological niches.

Here we test how the outcome of competition between *D. santomea* and *D. yakuba* varies across an ecologically relevant range of thermal environments. First, we study the distribution of *D. santomea* and *D. yakuba* on the island of São Tomé and find that temperature and seasonality differ between areas where these two species are found. We then manipulate temperature and the opportunity for competition in the lab to show that both species display maximal performance at moderate temperatures, but that at low temperatures *D. santomea* is able to outcompete *D. yakuba*, while the opposite is true at warmer temperatures. We then use a multigenerational ‘coexistence’ experiment to show that the relative abundances of *D. santomea* and *D. yakuba* are strongly affected by the thermal environment: at low temperatures, *D. santomea* and *D. yakuba* coexist, while at higher temperatures *D. yakuba* rapidly drives *D. santomea* to extinction. Our results show that the interaction between temperature and competition helps shape ecological differences between these two species; and, disturbingly, suggest that with warming temperatures, *D. yakuba* will be able to outcompete *D. santomea*, potentially contributing to extinction of this island endemic species.

## Materials and Methods

### The thermal niche of D. santomea and D. yakuba on São Tomé

We first qualitatively describe the thermal niche of *D. santomea* and *D. yakuba* found on the island of São Tomé using a previously published dataset of occurrence records for both species (Comeault et al. 2016) and climate data from the Worldclim database (bioclim variables 1-11; Table S1; http://www.worldclim.org/bioclim). We extracted values for bioclim variables 1-11 at a resolution of 2.5 arc-degrees, for each site, using the *raster* R library (Hijmans et al. 2019). Because the transect on São Tomé covers a relatively small geographic area – it spans only four unique sets of bioclim variables – we report the range of (1) mean annual temperatures (bioclim variable 1), (2) isothermality (bioclim variable 3: mean diurnal range of temperatures / annual range of temperatures), and (3) temperature seasonality (bioclim variable 4) for sites where either *D. santomea* or *D. yakuba* is the more abundant species, based on relative abundances reported in Figure S1 of Comeault et al. 2016, rather than summarize climatic variation using a typical decomposition-based method (e.g. principal components analysis). These three variables were chosen because mean annual temperature was strongly correlated with all ‘thermal’ bioclim variables (*r* > 0.9) except isothermality (*r* = 0.11) and seasonality (*r* = 0.36) and isothermality and seasonality showed more modest correlations with all other thermal bioclim variables (minimum *r* = 0.09, maximum *r* = 0.70; mean *r* = 0.26).

### Details of populations used for experiments

To measure performance under different temperatures and competitive environments we used individuals from genetically diverse laboratory populations of *D. santomea* and *D. yakuba*. We generated these populations by combining 5 male and 5 female offspring from each of 20 isofemale lines established from inseminated females collected on São Tomé. Females were collected between 1 and 14 February 2015 at the sites “lake7” (for *D. santomea*) and “monte7” (for *D. yakuba*) and the synthetic populations were created on 12 and 19 March 2015 after ~2-4 generations in the lab. The two resulting synthetic populations (D. *santomea*: “san_lake_7S” and *D. yakuba*: “yak_monte_7S”) were maintained at large population sizes spread over three to five 175 ml polypropylene bottles (Genesee Scientific, Morrisville, NC) for between 5 and 10 overlapping generations before initiating experiments.

### Temperature’s effect on competition between *D. santomea* and *D. yakuba*

We tested the relative performance of *D. santomea* and *D. yakuba* at each of three biologically relevant temperatures (18°C, 22°C, and 25°C) when maintained in isolation or together. To initiate this experiment, we placed 6 one to 9 day old female flies from the stock populations into individual 30 mL vials containing standard corn-starch medium. Sampling females from stock populations in this way results in >95% of the females being inseminated and actively laying viable eggs (see Supporting Information, Appendix I). To quantify performance in the absence of interspecific competition, 6 females of either *D. santomea* or *D. yakuba* were added to the vials. To quantify the effect of interspecific competition, three females of each species were added to the same vial. Therefore the total number of laying females remained constant between the “isolation” and “competition” treatments and adult flies and larvae experienced intraspecific competition only in the former treatment, while they experienced both intra and interspecific competition in the latter. We created a total of 30 replicate vials containing only *D. santomea* and only *D. yakuba*, and 30 “competition” replicates which contained both species. We then randomly assigned 10 replicates of each of the three resulting treatments – *D. santomea* in isolation, *D. yakuba* in isolation, or competition – to each of the three temperature treatments. Females were then allowed to lay eggs for 7 days and then removed from the vials. When removing the females we added a dampened (0.5% propionic acid) Kimwipe (Kimberly-Clark, catalog number: 34155) as a pupation site and to inhibit the growth of fungi. As a measure of performance across temperatures and in different competitive environments we counted the total number of each sex of adult fly that ecolsed from each vial over the following 23 days. While we counted the total number of flies, we focus analyses below on male counts because male *D. santomea* and *D. yakuba* are easily and unambiguously identified based on pigmentation: *D. santomea* males are a solid yellow and *D. yakuba* males have black pigmentation on the last three tergites of their abdomen (females show some variation in color). Despite potential for misclassification of females, we found no difference in sex ratio across treatments or temperatures (binomial GLM; Figure S1) and summarize results from analyzing all offspring in the Results and Figure S2. We stopped counting emerging flies after 23 days because this duration spanned peak eclosion at all temperatures (Figure S3).

To test whether temperature affected the outcome of competition between *D. santomea* and *D. yakuba* we first modeled the mean (per-female) number of male flies that eclosed from an experimental replicate as a function of temperature, competition treatment (three levels: *D. santomea*, *D. yakuba*, or interspecific competition), and the interaction between temperature and competition. This model was fit using the *glm* function in R (Core Team 2017) assuming Poisson distributed error. To determine if there was a significant interaction between temperature and competition treatment, we conducted a likelihood ratio test (LRT) that compared the model described above to one that lacked the interaction term using the *anova* function in R. Because this analysis identified a significant interaction between temperature and competition (see Results) on the mean number of male offspring produced, we also fit independent generalized linear models, splitting the data by temperature treatment. We then used Tukey’s post-hoc tests, as implemented with the *glht* function in the *multcomp* R package (Hothorn et al. 2008), to test for significant pairwise differences in performance between competition treatments, for each temperature treatment. Finally, for replicates that contained both *D. santomea* and *D. yakuba* (i.e. the “competition” treatment) we tested for differences in performance between these species, at different temperatures, by conducting dependent-samples sign-tests (these data are naturally paired by replicate) as implemented by the *SIGN.test* function in the *bsda* R package (Arnholt and Evans 2017). Because *D. santomea* on the island of São Tomé are found in cooler environments than *D. yakuba*, we predicted that *D. santomea* would outcompete *D. yakuba* at 18°C and *D. yakuba* would outcompete *D. santomea* at 22°C and 25°C.

### Competition during experimental sympatry at different temperatures

We found a significant interaction between temperature and competition treatments in the single-generation experiment described above. Building on this result, we next tested whether competition between *D. santomea* and *D. yakuba* at different temperatures could lead to one species competitively displacing the other. To test this, we created two-species experimental ‘communities’ in mesh cages (24.5 cm × 24.5 cm × 24.5 cm; www.bugdorm.com) and maintained them at either 18°C or 25°C (four replicates at each temperature). Each cage was prepared by adding ~1 inch of dampened coconut fiber as a substrate to help maintain a relative humidity between ~40 and 80%. We then added two 175 ml polypropylene bottles (Genesee Scientific, Morrisville, NC) containing ~30ml of standard cornmeal medium and a damped kimwipe to each cage along with 24 males and 24 females of both *D. santomea* and *D. yakuba* (96 flies founded each experimental community). The flies used to initiate this experiment were collected 14 hours after eclosion and were 3 days old when introduced to the cages. Every two weeks, we added two fresh bottles containing cornmeal medium and bottles were removed after they were in the cages for at least four weeks (see Table S2 for details). We then randomly sampled flies from each experimental cage at 34 days, 57 days, 90 days, 113 days, and 148 days after initiating the experiment. 148 days is ~10 generations at 18°C and ~12 generations at 25°C for both of these species. To test for consistent changes in the relative abundance of the two species within each cage we conducted Mann-Kendall Trend Tests on the proportion of *D. santomea* within each cage. We expect a significant (and consistent) change in the proportion of *D. santomea* within each cage if it was either at a competitive advantage (increase) or disadvantage (decrease) over *D. yakuba*.

## Results

### The thermal niche of D. santomea and D. yakuba on São Tomé

We found that along the altitudinal transect on São Tomé, *D. santomea* is more abundant than *D. yakuba* at cooler sites, where mean annual temperatures range from 17.8 to 20.6°C and seasonality (in units of standard deviations in temperature) ranges from 8.24 to 8.29 (Figure 1). By contrast, *D. yakuba* is more abundant at sites experiencing higher mean annual temperatures (20.6 to 25.5°C) and stronger seasonality (8.27 to 8.35; Figure 1). Isothermality – measured as the mean diurnal range over the annual temperature range – was effectively invariable across the sampled sites (range = 0.70 to 0.71). These qualitative patterns are consistent with previously published results that suggest that *D. santomea* has a behavioral preference for cooler temperatures than *D. yakuba* (Matute et al. 2009).

**Figure 1.**
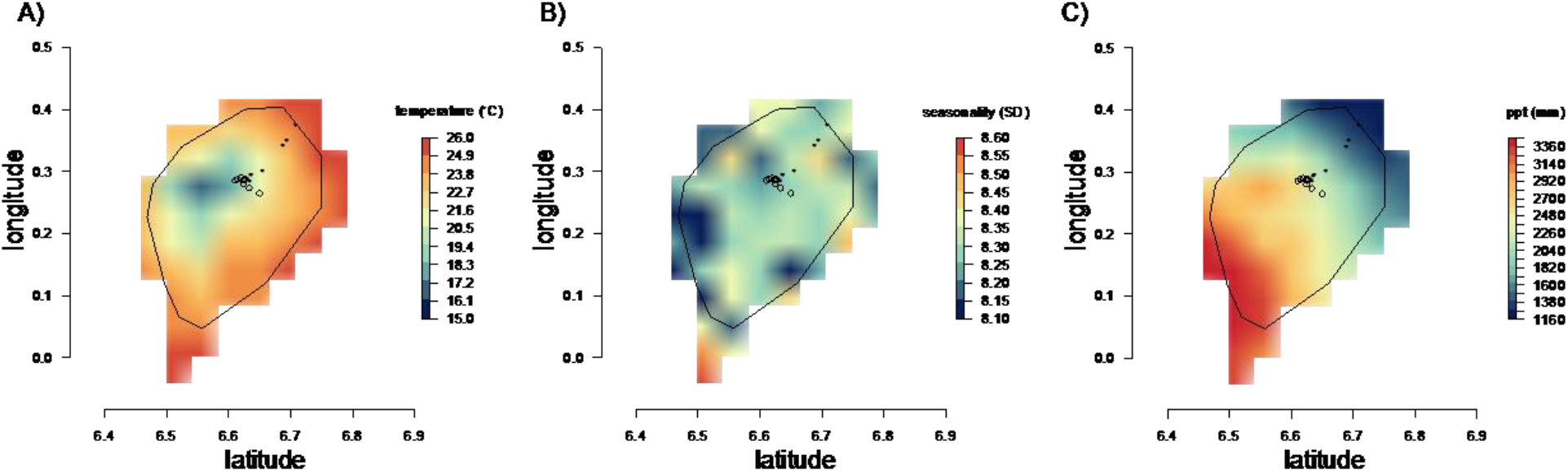
Mean annual temperature (A) and temperature seasonality (B) and mean annual precipitation (C) on the island of São Tomé. Black line outlines the coast of São Tomé, white space is the Atlantic Ocean. Filled black points denote sites where *D. yakuba* is the more abundant species and open black circles represent sites where *D. santomea* was more abundant. SD = standard deviation.

### Temperature’s effect on competition between D. santomea and D. yakuba

When raised in isolation, performance, as measured as the number of emergent male offspring, varied with temperature for both *D. santomea* and *D. yakuba* (GLMs: *D. santomea*: X_2_ = 14.26; *P* = 0.0008; *D. yakuba*: X_2_ = 8.54; P = 0.014). *Drosophila santomea* produced significantly more male offspring when maintained at 22°C (mean = 8.1; S.D. = 0.5) compared to when they were maintained at 18°C (mean = 4.8; S.D. = 0.3) (Tukey’s HSD: *Z* = 2.5; *P* = 0.03) or 25°C (mean = 4.5; S.D. = 0.3) (Tukey’s HSD: *Z* = 3.6; *P* = 0.001). *Drosophila yakuba* also produced significantly more male offspring when maintained at 22°C (mean = 8.0; S.D. = 0.5) compared to 25°C (mean = 5.1; S.D. = 0.3) (Tukey’s HSD: *Z* = 2.9; *P* = 0.012), but there was not a significant difference (Tukey’s HSD: *P* > 0.1) in performance between either 22°C and 18°C (mean = 5.9; S.D. = 0.4) or 25°C and 18°C. These results suggest that both *D. santomea* and *D. yakuba* perform best at temperatures in the low 20’s (°C), consistent with previous results (Matute et al. 2009).

In addition to temperature, the interaction between temperature and competitive environment had a strong effect on performance (LRT: X_2_ = −28.69; *P* < 1.0 × 10_−4_; Figure 2A). When maintained together (i.e., with the opportunity for interspecific competition) at 18°C, *D. santomea* had higher performance than *D. yakuba* (Tukey’s HSD: *Z* = 3.09; *P* = 0.01) and *D. yakuba* performed worse than when maintained in isolation (Tukey’s HSD: *Z* = 2.65; *P* = 0.04) (Figure 2A). When maintained together at 25°C, *D. yakuba* outperformed *D. santomea* (Tukey’s HSD: *Z* = 3.55; *P* = 0.002), but there was not a significant difference in *D. santomea*’s performance when maintained in isolation or together with *D. yakuba* at 25°C (Tukey’s HSD: *Z* = 1.42; *P* = 0.48). At 22°C, there was no significant effect of competitive environment on per-female performance in either *D. santomea* or *D. yakuba* (Tukey’s HSD: all *P* > 0.05). We report results from parallel analyses including both female and male offspring in Appendix II and Figure S1 (Supporting Information). The only difference we observed when analyzing total offspring (i.e. male and female offspring both included) was that *D. yakuba that* experience interspecific competition at 22°C produce fewer offspring than when maintained in the absence of interspecific competition at 22°C (Tukey’s HSD: *Z* = −2.724; *P* = 0.0325; Figure S2).

**Figure 2.**
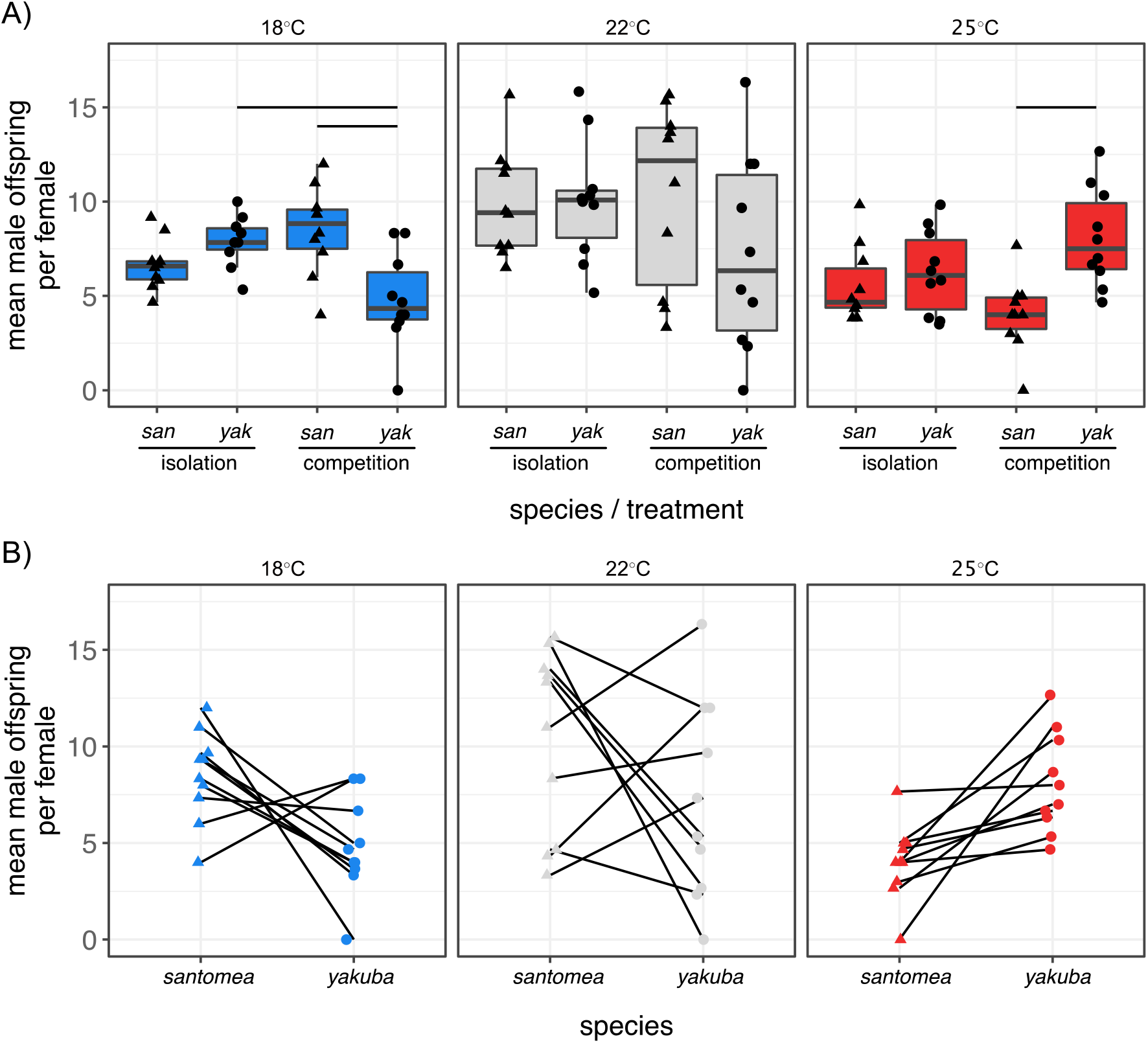
The effect of temperature and competition on performance. A) The mean number of offspring per female when *D. santomea* (*san*) and *D. yakuba* (*yak*) were raised with the opportunity for intraspecific competition only (“isolation”) or with both intra and interspecific competition (“competition”). Groups that showed a significant difference in the mean number of offspring produced are indicated by horizontal bars (Tukey’s pairwise contracts; generalized linear models run separately for each temperature). When maintained at 18°C and in competition with *D. santomea*, *D. yakuba* showed a significant reduction in fitness (leftmost panel of A), while the opposite is true at 25°C (rightmost panels). At 22°C, there is a large variance in the number of offspring that emerged and we did not detect a difference between *D. santomea* and *D. yakuba*. B) Competition data presented in A), but as paired data. Lines connect the number of male *D. santomea* and *D. yakuba* for each of 10 replicates conducted at each temperature.

When treating performance of *D. santomea* and *D. yakuba* within each replicate of the interspecific competition experiment as paired, *D. santomea* tended to produce more offspring than *D. yakuba* at the low temperature (18°C), resulting in a marginally significant effect (*S* = 8, *P* = 0.055; Figure 2B, left panel), *D. yakuba* produced more offspring than *D. santomea* at the high temperature (22°C; *S* = 0, *P* = 0.001; Figure 2B, right panel), and there was no consistent difference in the number of offspring produced by either species at 22°C (*S* = 9, *P* = 0.828; Figure 2B, middle panel).

### Competition during experimental sympatry at different temperatures

Consistent with the results we observed over the course of a single generation, when maintained together at 25°C the proportion of *D. santomea* monotonically decreased over the course of the experiment (*S* = −9, −9, and −12; *P* = 0.036, 0.036, and 0.013, respectively, for the three cages that maintained viable populations over the entire course of the experiment). Under our experimental conditions, *D. santomea* went extinct in three of the four cages maintained at 25°C after 57 days (4-5 generations; in the fourth cage only one of 67 sampled individuals was *D. santomea* (Figure 3 & S4)). By contrast, when maintained together at 18°C, the proportion of *D. santomea* in each cage did not monotonically change over the course of the experiment (*S* = 7, 3, 5, and −7; *P* = 0.13, 0.35, 0.23, and 0.87, respectively), and *D. santomea* was frequently more abundant than *D. yakuba* in cages maintained at 18°C (Figure 3). This trend of *D. santomea* being more abundant than *D. yakuba* in cages maintained at 18°C was not statistically significant (generalized linear mixed model assuming Poisson distributed error with sample date and cage as random effects: fixed effect of species: *Z* = −1.314; *P* = 0.19).

**Figure 3.**
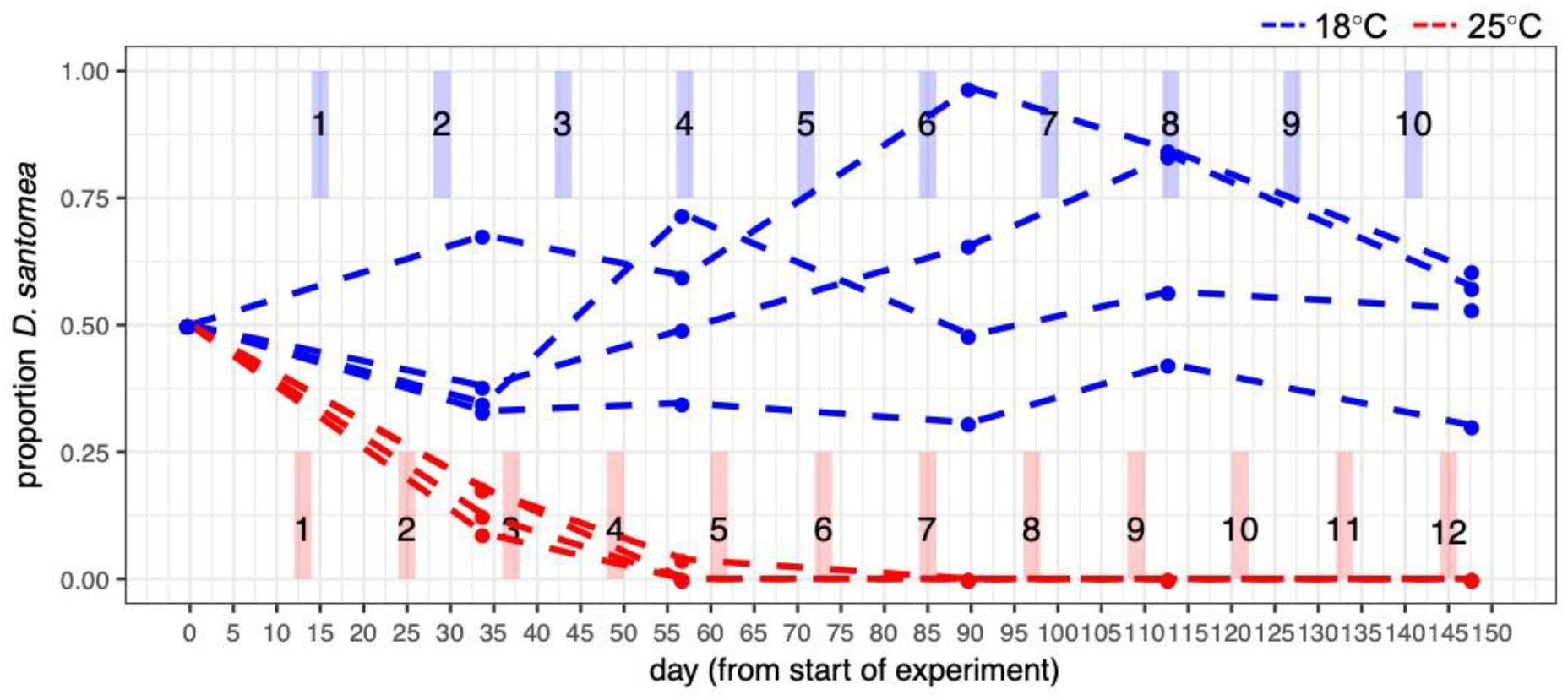
The multi-generation effect of temperature on the coexistence of *D. santomea* and *D. yakuba* in experimental sympatry. When *D. santomea* and *D. yakuba* are maintained together in cages at 18°C (blue lines) the proportion of *D. santomea* within the cages tends to be near or above 0.5. By contrast, when maintained at 25°C (red lines), *D. santomea* is competitively excluded from the cages within ~4-7 generations. The approximate number of generations is indicated, for both temperature treatments, by shaded rectangles (blue = generation times when maintained at 18°C; red = generation times when maintained at 25°C).

## Discussion

Our results show how the thermal environment modifies competitive outcomes between the co-occurring sibling species *D. yakuba* and *D. santomea*. We find that in the absence of interspecific competition, both species display the highest performance at a moderate temperature of ~22°C (Figure 2A). However, the competitive dominance hierarchy displayed by these two species depends on the thermal environment under which competition takes place: at higher temperatures *D. yakuba* outcompetes *D. santomea*, while at lower temperatures *D. santomea* outcompetes *D. yakuba* (Figure 2A). The direction of competitive advantage therefore depends on the thermal environment under which competition takes place and is consistent with the thermal niche these two species inhabit in nature (Figure 1). These results suggest that the temperature-dependent outcome of competition between these two species is contributing to the maintenance of their narrow band of sympatry at mid elevations on the island of São Tomé.

### Competition’s role in moderating local adaptation to different thermal environments

We did not observe a symmetrical effect of competition at the low and high temperatures that we tested, as *D. santomea* did not competitively exclude *D. yakuba* within ~10 generations of being maintained together at 18°C (Figure 3). While *D. santomea*’s competitive advantage at lower temperatures meant that it tended to be the more abundant species in experimental enclosures maintained at 18°C, this effect was minor (see blue lines in Figure 3 and Figure S4). Given *D. yakuba* is able to maintain high performance at relatively low temperatures, even in the presence of *D. santomea*, what stops *D. yakuba* from moving to higher elevation habitats on São Tomé? One variable that our experiments do not account for is the behavioral preferences of *D. santomea* and *D. yakuba* for different environments. Laboratory experiments have shown that *D. santomea* displays a behavioral preference for cooler environments than *D. yakuba* (Matute et al. 2009). *D. yakuba* is also a broadly distributed generalist species found across sub saharan Africa (Lachaise et al. 1988; Yassin et al. 2016), while *D. santomea* has primarily been found in association with figs and is endemic to São Tomé (Lachaise et al. 1988, 2000). It is therefore likely that multiple ecological factors, including temperature, humidity, habitat type, and diet, affect the realized distributions of *D. santomea* and *D. yakuba* on São Tomé. Future studies testing competition and performance on different diets and at temperatures below 18°C (the lowest we tested) are needed to more fully understand the factors defining the ranges of *D. santomea* and *D. yakuba*. Our results show that differences in competitive ability across thermal environments is one factor that is likely to affect the realized niches of these two species.

### Context dependent responses to climate change

Climate change exposes species to warmer mean annual temperatures and their demographic and evolutionary responses to warmer temperatures will depend on both direct and indirect effects of the thermal environment. For example, species may show direct responses to different thermal environments through the evolution of novel or different physiological traits (Eliason et al. 2011; Cooper et al. 2012; Campbell-Staton et al. 2020). In the context of climate change, a species will only be able to persist and maintain demographically viable populations if the benefit of direct evolutionary responses to warmer (or more variable) temperatures is not outweighed by negative changes in biotic interactions. Previous studies have, for example, shown how changes in the thermal environment and / or the community of interspecific competitors can lead to lower fitness or rapid extirpation (Davis et al. 1998*a*; Alexander et al. 2015). Competitive exclusion has therefore been discussed as an important outcome of climate change in a number of species (Finstad et al. 2011; Bulgarella et al. 2014), including even between anatomically modern humans and Neaderthals (Banks et al. 2008).

One of the predictions generated from our results with respect to competition between *D. santomea* and *D. yakuba* is that under warmer mean daily temperatures, the highland forest endemic species, *D. santomea*, may be challenged by competitive exclusion by *D. yakuba*, ultimately leading to extinction. There are three caveats to consider when interpreting this prediction. First, the multigenerational experiment we conducted that showed competitive exclusion of *D. santomea* by *D. yakuba* at 25°C (Figure 3) did not include appropriate experimental controls, where the two species were maintained in isolation. We therefore assume that *D. santomea* is capable of maintaining viable populations at 25°C under the same experimental conditions in the absence of competition. Two lines of evidence support this assumption. First, when we raised *D. santomea* in isolation at 25°C, over a single generation, each female produced an average of 5.6 male and 6.5 female offspring (Figure 2), suggesting they can maintain positive population growth at this temperature. Second, previous work on thermal performance traits in *D. yakuba* and *D. santomea* show that males and females of both species remain fertile and reproductively active at 24°C (Matute et al. 2009). We therefore interpret the multigenerational experiment summarized in Figure 3 as providing evidence for competitive exclusion of *D. santomea* at 25°C. Under this interpretation, this result highlights the importance of considering both direct and indirect effects when estimating the impacts that climate change will have on biodiversity.

Second, we did not account for frequency- or density-dependent processes that may alter competitive advantages over the course of the multigenerational experiment. For example, if *D. santomea* exhibited strong competitive dominance over *D. yakuba* at 18°C one prediction is that they would consistently be the more abundant species across the course of the experiment. While there was a weak trend in our data to suggest *D. santomea* tended to be more abundant than *D. yakuba* at 18°C, this trend was not statistically significant (GLMM: *P* = 0.19; Figure S4). Our method of estimating relative abundances within experimental cages (non-standardized sampling effort across cages and sampling dates) also did not allow us to test density-dependent effects. Future future work is needed to understand the density-dependence of competitive outcomes between *D. santomea* and *D. yakuba* across different thermal environments.

Third, we did not test whether local adaptation to the experimental environment (biotic and abiotic) altered the competitive interaction between *D. santomea* and *D. yakuba*. Studies of local adaptation in these and other species of Drosophilid flies have shown that they can show evolutionary responses to selection in as few as 10 generations (Matute 2010a, 2010b; Bergland et al. 2014; Tobler et al. 2014; Comeault et al. 2016; Behrman et al. 2018). Local adaptation in *D. santomea*, *D. yakuba*, or both species therefore could have altered the nature of competition or population growth rates over the course of our experiment. Future experiments that either control for evolution or explicitly measure evolution’s effects on species interactions under different environmental conditions have the potential to greatly increase our understanding of species’ responses to environmental change (e.g. (Germain et al. 2020)).

In addition to competitive interactions, temperature is likely to affect, and modify, other types of biotic interactions. For example, the fitness cost of parasites can vary with temperature in a similar way to competition (Lashomb et al. 1987; Vale et al. 2008) and, in some cases, mutualists can become parasites when temperature increases (Baker et al. 2018). In other cases, symbionts can alter the thermal tolerance of their host and act to facilitate colonization of new niches (Russell and Moran 2006). Together with these examples, the results we present here illustrate how predicting a species’ response to environmental change (such as global warming) requires an understanding of both the direct and indirect effects imposed by that change.

### A role for biotic interactions in range-shifts associated with climate change

An often discussed (and observed) response that species have to climate change is a poleward or upslope shift in their range (Parmesan 2006; Colwell et al. 2008; Lenoir et al. 2008; Schuetz et al. 2019). Range-shifts can be driven by a species tracking favorable abiotic conditions, such as temperature, but the rate and extent of range-shifts is likely to vary among species, resulting in “community reorganization” (Van der Putten 2012). Climate associated range-shifts are therefore likely to affect interspecific interactions in at least two ways. First, they can change the identity of the interacting members of a community and, in the case of interspecific competition, alter competitive dominance hierarchies (Alexander et al. 2015). Second, they can change the environmental context of the interaction – such as when tracking one environmental variable results in a change in a second – and alter the outcome of specific interactions (Bronstein 1994; Davis et al. 1998*a*; Tylianakis et al. 2008; Chamberlain et al. 2014; Harrower and Gilbert 2018). The results we have presented here provide an example of the latter and point to the importance of quantifying interspecific interactions under different environments in order to better predict responses to climate change. This may be particularly important for tropical endemic species threatened by the invasion of sibling species that can displace them through competitive exclusion.

## Acknowledgments

We thank D. Turrisini, L. Sliger, and B. Cooper for help in the field, and the Matute lab, Bangor University’s “Journal Club in the Pub”, and three anonymous reviewers for constructive feedback on previous drafts. This work was supported by the National Science Foundation (Dimensions of Biodiversity award number 1737752 to DRM). The funders had no role in any aspect of study design, data collection and analysis, or decisions with respect to publication.

