## Supporting Information for "Temperature-dependent competitive outcomes between the fruit flies *Drosophila santomea* and *D. yakuba*"

This file contains:

**Appendix I**: The frequency of fertility in females sampled directly from population bottles.

**Appendix II**: Results when analyzing both male and female offspring together.

**Table S1**: Description of bioclimatic variables used to describe the thermal environments *D. santomea* and *D. yakuba* inhabit on São Tomé.

**Table S2**: Maintenance schedule and sampling of the two-species community cages.

**Figure S1**: Offspring sex ratios across competition and temperature treatments.

**Figure S2**: Interaction between temperature and competitive environment on the total number of offspring produced per female.

**Figure S3**: Cumulative eclosion curves for *D. santomea* and *D. yakuba* at different temperatures.

**Figure S4**: Abundances of *D. santomea* and *D. yakuba* males sampled over the course of the multigeneration "sympatry" experimental cages.

**Appendix I:** The frequency of fertility in females sampled directly from population bottles.

To determine the frequency of inseminated female flies with the stock bottles used to initiate the experiment described in the section "*Temperature's effect on competition between* D. santomea *and* D. yakuba" of the main text we collected 140 females (70 females of each species) and isolated them in individual vials containing cornmeal medium. As for the main experiment reported in text, we collected females that were between one and 9 days old from bottles. Of the 140 vials containing females only 4 did not contain offspring (larvae, pupae, and/or adult progeny) after 14 days. Three of these four were vials initiated with female *D. santomea* and one with *D. yakuba*.

**Appendix II:** Results when analyzing both male and female offspring together.

When analyzing the mean total number of offspring produced by each female (i.e. counting both male and female offspring), as with the results reported in the main text, the interaction between temperature and competitive environment had a strong effect on performance (LRT: $X_2$ = -65.317; $P < 0.001$; Figure S2). In the presence of interspecific competition at 18ºC, *D. santomea* had higher performance than *D. yakuba* (Tukey's HSD: $Z$ = -4.130; $P < 0.001$) and *D. yakuba* performed worse than when maintained in isolation (Tukey's HSD: $Z$ = -4.130; $P < 0.001$) (Figure S2). In the presence of interspecific competition at 25ºC, *D. yakuba* outperformed *D. santomea* (Tukey's HSD: $Z$ = 6.076; $P < 0.001$), but there was not a significant difference in *D. santomea*'s performance when maintained in isolation or together with *D. yakuba* at 25ºC (Tukey's HSD: $Z$ = 1.869; $P$ = 0.2402). At 22ºC, the only significant difference in per-female performance was between *D. yakuba* females in isolation and *D. yakuba* experiencing interspecific competition (Tukey's HSD: $Z$ = -2.724; $P$ = 0.0325), with interspecific competition tending to reduce the mean number of female offspring produced at this temperature (Figure S2).

**Table S1**. Description of bioclimatic variables used to describe the thermal *environments D. santomea* and *D. yakuba* inhabit on São Tomé.

| Bioclim variable number | Description |
|---|---|
| 1 | Annual mean temperature |
| 2 | Mean Diurnal Range (Mean of monthly (max temp - min temp)) |
| 3 | Isothermality (BIO2/BIO7) (*100) |
| 4 | Temp Seasonality (standard deviation *100) |
| 5 | Max Temp of Warmest Month |
| 6 | Min Temp of Coldest Month |
| 7 | Temp Annual Range (BIO5-BIO6) |
| 8 | Mean Temp of Wettest Quarter |
| 9 | Mean Temp of Warmest Quarter |
| 10 | Mean Temp of Driest Quarter |
| 11 | Mean Temp of Coldest Quarter |

**Table S2.** Maintenance schedule and sampling of the two-species community cages.

| Date | Notes |
|---|---|
| 22 Oct 2015 | - START -<br>- founding flies added to each two-species community cage<br>- 24 males and 24 females of each species (72 hours after eclosion).<br>- RH% (temp) = 80 (25ºC), 50 (18ºC) |
| 02 Nov 2015 | - RH = 54 (25ºC), 70 (18ºC) |
| 03 Nov 2015 | - RH = 82 (25ºC), 73 (18ºC) |
| 05 Nov 2015 | - added 2 fresh bottles of food to cages |
| 09 Nov 2015 | - RH = 80 (25ºC), 40 (18ºC) |
| 25 Nov 2015 | - sampled cages<br>- added 2 fresh bottles of food to cages<br>- removed 2 oldest bottles |
| 17 Dec 2015 | - added 2 fresh bottles to 25 degree cages<br>- removed 2 oldest bottles |
| 18 Dec 2015 | - sampled cages<br>- added 2 fresh bottles to 18 degree cages<br>- removed 2 oldest bottles |
| 4 Jan 2016 | - added 2 fresh bottles to 25 degree cages<br>- removed 2 oldest bottles<br>- !! one of the cages at 25 degrees went extinct !! |
| 20 Jan 2016 | - sampled cages<br>- added 2 fresh bottles to 25C and 18C cages<br>- removed 2 oldest bottles |
| 12 Feb 2016 | - sampled cages<br>- added 2 fresh bottles to 25C and 18C cages<br>- removed 2 oldest bottles |
| 18 Mar 2016 | - sampled cages<br>- END - |

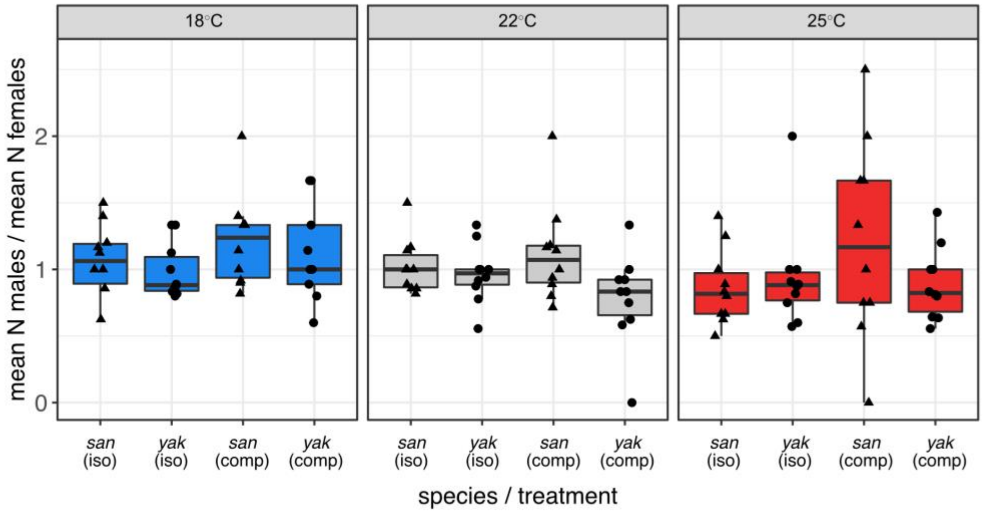


**Figure S1.** Offspring sex ratios across competition and temperature treatments for *D. santomea* (*san*) and *D. yakuba* (*yak*). We found no effect of competition treatment (levels: isolation, competition), temperature (levels: 18°C, 22°C, 25°C), or the interaction between competition and temperature on the ratio of males to females across replicates. Effects were tested with nested binomial generalized linear models and likelihood ratio tests (all $P > 0.1$).

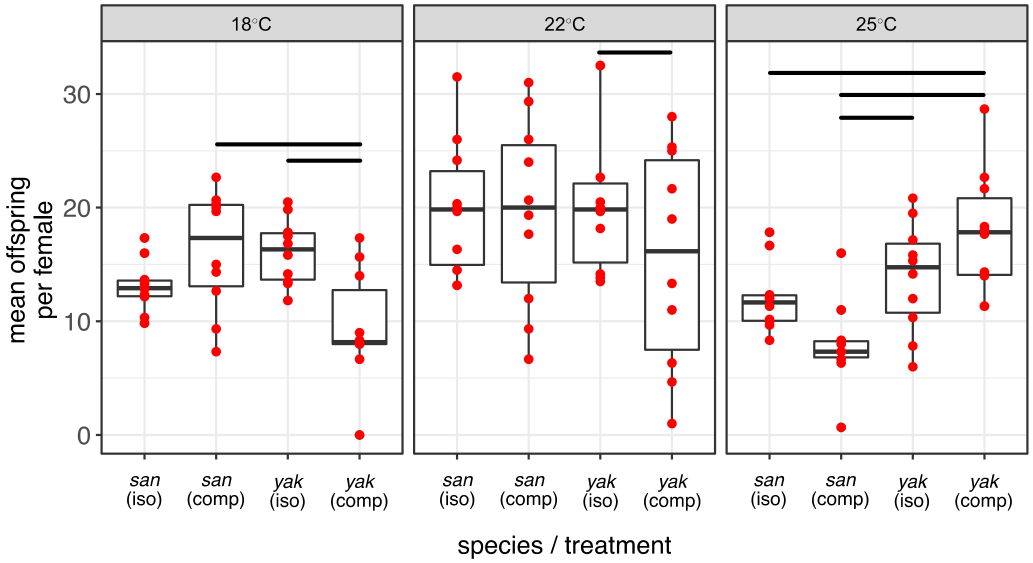


**Figure S2.** The effect of temperature and competition on mean offspring produced per female. The number of offspring is summarized for each species when raised either in the absence of interspecific competition (iso) or experiencing interspecific competition (comp). Groups that showed a significant difference in the mean number of offspring produced are indicated by horizontal bars (Tukey's pairwise contracts of generalized linear models run separately for each temperature).

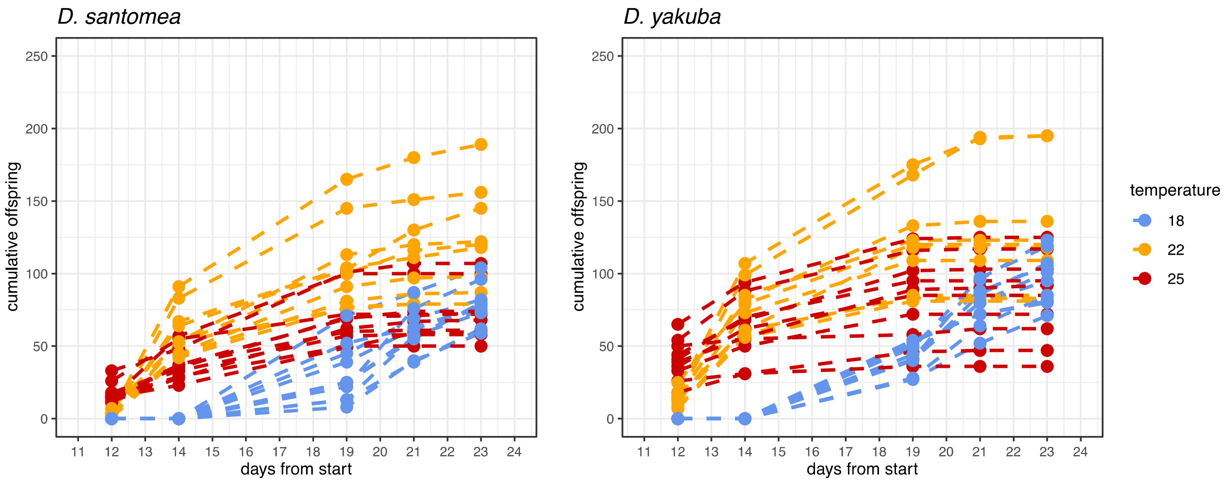


**Figure S3**. Cumulative eclosion curves for *D. santomea* (left panel) and *D. yakuba* (right panel) when raised at each of the three temperatures used in experiments reported in the main text. Temperature is in degrees centigrade.

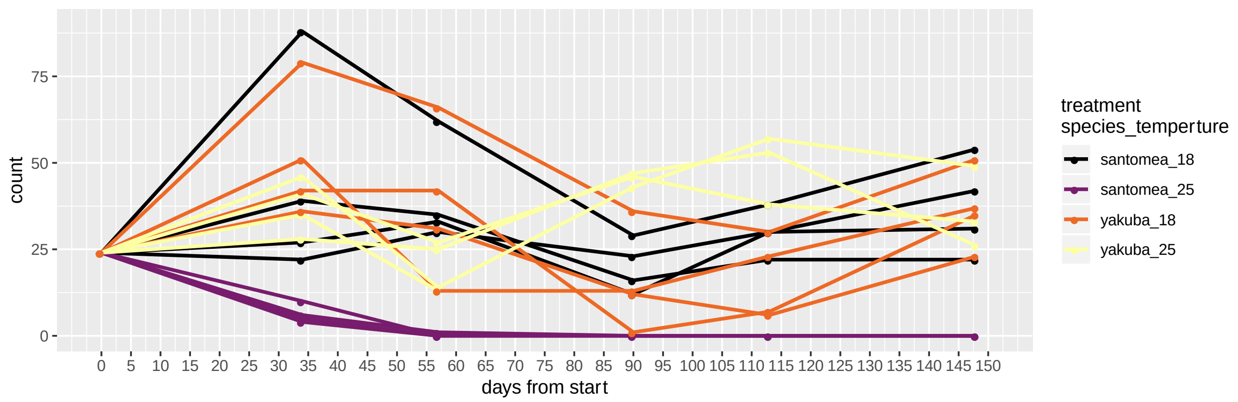


**Figure S4**. Abundances (counts) of *D. santomea* and *D. yakuba* over the course of the multigenerational “coexistence” experiment. Counts are of male individuals only. See Figure 3 of the main text for results presented as proportions. Note that counts are derived from samples taken from within each experimental cage and should not be used as an estimate of total population size as sampling effort was not rigorously standardized when sampling cages (e.g. a cage with a high density of flies was not sampled for the same amount of time as one with a low density of flies).
